# Microhabitat partitioning correlates with opsin gene expression in coral reef cardinalfishes (Apogonidae)

**DOI:** 10.1101/744011

**Authors:** Martin Luehrmann, Fabio Cortesi, Karen L. Cheney, Fanny de Busserolles, N. Justin Marshall

## Abstract

1. Vertebrates exhibit diverse visual systems that vary in terms of morphology, number and distribution of spectrally distinct photoreceptor types, visual opsin genes and gene expression levels.
2. In fish, such adaptations are driven by two main factors: differences in the light environment and behavioural tasks, including foraging, predator avoidance and mate selection. Whether visual systems also adapt to small-scale spectral differences in light, between microhabitats, is less clear.
3. We suggest that differences in microhabitat use by cardinalfishes (Apogonidae) on coral reefs drive morphological and molecular adaptations in their visual systems. To test this, we investigated diurnal microhabitat use in 17 cardinalfish species and assessed whether this correlated with differences in visual opsin gene expression and eye morphology.
4. We found that cardinalfishes display six types of partitioning behaviours during the day, ranging from specialists found exclusively in the water column to species that are always hidden inside the reef matrix.
5. Using data on visual opsin gene expression previously characterized in this family, it was discovered that species in exposed habitats had increased expression of the short-wavelength sensitive violet opsin (*SWS2B*) and decreased expression of the dim-light active rod opsin (*RH1*). Species of intermediate exposure, on the other hand, expressed opsins that are mostly sensitive to the blue-green central part of the light spectrum (*SWS2As* and *RH2s*), while fishes entirely hidden in the reef substrate had an increased expression of the long-wavelength sensitive red opsin (*LWS*).
6. We found that eye size relative to body size significantly differed between cardinalfish species, and relative eye size decreased with an increase in habitat exposure.
7. Retinal topography did not show co-adaptation with microhabitat use, but instead with feeding mode.
8. We suggest that, although most cardinalfishes are nocturnal foragers, their visual systems are also adapted to both the light intensity and the light spectrum of their preferred diurnal microhabitat.

## Introduction

Animal visual systems are functionally diverse, with differences at the morphological and the molecular level. In fish, this diversity is mainly driven by differences in the availability of light (Hauser and Chang, 2017; Land and Nilsson, 2002), but can also be due to differences in habitat complexity (Collin and Shand, 2003; Hughes, 1977) or specific behavioural tasks, e.g. foraging or sexual selection (reviewed in Hauser and Chang, 2017; Price, 2017). In aquatic environments, differences in light environments arise from wavelength selective light absorption and scattering due to depth and various sizes of particles (Lythgoe, 1979). For example, the deep-sea has a blue-shifted light environment and consequently, deep-sea species generally possess photoreceptors that are maximally sensitive to blue light (~ 480 nm) (Partridge et al., 1992).

Morphologically, visual systems may differ in eye size, shape, and at the retinal level in functional type, number and/or distribution of neural cells including photoreceptors. Morphological changes to boost sensitivity in low-light conditions, for example, may include rod-dominated retinas, increased relative eye size, or a higher photoreceptor-to-ganglion cell summation ratio (de Busserolles and Marshall, 2017; Kelber and Roth, 2006; Warrant, 2004).

At the molecular level, changes in the visual opsins and associated light-sensitive chromophores may also reflect functional adaptation by shifting photopigment spectral sensitivity (Hunt and Collin, 2014). Photopigments are comprised of opsins, membrane-bound proteins with G-protein coupled receptor function to which a vitamin A-derived chromophore is covalently bound. The specific combination of opsin and chromophore determines to what part of the electromagnetic spectrum the photopigment is maximally sensitive to (λ_max_). Opsins are classified according to their λ_max_ values, their phylogeny, and their photoreceptor specificity (Hunt et al., 2014). Percomorph fishes have multiple opsin types, including a rod-specific opsin (rhodopsin, RH1) used for scotopic vision, and various cone-specific opsins that are used for photopic and colour vision. These are the short-wavelength-sensitive cone opsins expressed in single cones: SWS1 (UV), SWS2B (violet), SWS2Aα and SWS2Aβ (blue); and the mid- to long-wavelength-sensitive cone opsins expressed in double cones: RH2B (blue-green), RH2A (green), and LWS (yellow/red) (Cortesi et al., 2015; Hunt and Collin, 2014).

The λ_max_ of a photoreceptor depends primarily on i) variations in the amino acid sequence of the opsin, ii) the type of associated chromophore (Vitamin A1 or A2), and iii) the levels of expression of the different opsin genes (reviewed in Carleton et al., 2016). The suite of spectral sensitivities a fish possesses at any one time may also be plastic, with adaptations to changing visual demands over environmental and/or ontogenetic, or other time scales observed in several species (reviewed in Carleton et al., 2016; Marshall et al., 2018).

Visual systems adapt to large-scale lighting differences due to habitat depth, season or type (Lythgoe, 1979; Lythgoe et al., 1994; Muntz, 1982). In addition, it is hypothesized that fish vision may also be tuned to smaller-scale differences in light - between microhabitats (Lythgoe, 1979; Marshall et al., 2003). This idea, however, has not been tested rigorously (Cummings and Partridge, 2001; Sabbah et al., 2011). It remains to be tested, for example, whether this phenomenon may contribute to visual system diversification among fishes living on coral reefs, one of the most diverse ecosystems on earth, where habitat partitioning is particularly common (reviewed in Williams, 1991).

Here, in order to control for potentially confounding factors like phylogenetic constraint, we focused on a group of closely related reef fishes with remarkable visual system diversity, the cardinalfishes (Apogonidae) (Fishelson et al., 2004; Luehrmann et al., 2019). These fishes are common on shallow tropical coral reefs, are one of the most abundant reef fish families, and are predominantly nocturnal foragers (Marnane and Bellwood, 2002). During the day, they aggregate in large multi-species groups in and around coral heads (Gardiner and Jones, 2005; Greenfield and Johnson, 1990) where they carry out social behaviours, such as pair formation and mating (Kuwamura, 1983; Kuwamura, 1985; Saravanan et al., 2013). A previous survey of seven species found that in these multi-species aggregations fish display strict microhabitat partitioning among the same diurnal refuge sites, with some species found predominantly outside, and others within or below coral structures (Gardiner, 2010).

To test whether their visual system design is related to microhabitat use, we compared the microhabitat partitioning behaviour of 17 cardinalfish species to morphological and molecular differences in their visual systems. First, we conducted an ecological assessment of habitat partitioning in these focal species. Second, we tested whether their opsin gene expression, and/or relative eye size – as a proxy for light sensitivity (Land, 1990), correlated with diurnal microhabitat use. We used our previous opsin expression data which showed that cardinalfishes express multiple visual opsins, and that based on differences in opsin gene expression and spectral sensitivities, species can be placed into five, possibly functionally distinct, groups (Luehrmann et al., 2019). Third, the retinal photoreceptor and/or ganglion cell topographies of five cardinalfish species from different microhabitats were determined to gain additional insight into these fishes’ adaptations to their environment. Finally, visual system diversity was also tested for correlation with cardinalfish feeding ecology and activity period.

## Materials and Methods

### Microhabitat use assessment

Underwater visual surveys were conducted on SCUBA to determine microhabitat use of 23 cardinalfish species (Fig. 1a, Table 1) on reefs surrounding Lizard Island (14° 40’S, 145° 28’E), Great Barrier Reef, Australia. Fish counts were conducted between 6.30 am and 4 pm, from 3-14 March 2017. Data for *Apogonichthyoides melas* and *Pterapogon cf. mirifica* was taken from counts between 10 Feb. and 20 April 2015, as these species were not found during counts in 2017. Counts were performed as spot-counts at 111 sites distributed over eight different locations, with a site defined as a separate coral head, outcrop or boulder located > 5 meters apart (Fig. 1b). When a site was encountered, we approached slowly and waited for several minutes to ensure fish behaviour was not disturbed. Then, we recorded individual animal numbers to 20 and estimated larger groups to the nearest 50. We avoided double counting of sites by navigating around the locations systematically. We counted fish in four distinct microhabitat partitions defined as per Fig. 1c. For microhabitat partitioning analysis, we only used species for which at least ten individuals were counted at > 3 different sites, and then calculated the frequency of occurrence at each microhabitat partition as a proportion of total individuals counted per species (Fig. S1, Table 1). To identify patterns of microhabitat use, we then used hierarchical cluster analysis [Ward.D2, bootstrap=100, R-package: pvclust (R Core team, 2014; Suzuki and Shimodaira, 2006)] (Fig. 2).

**FIGURE 1.**
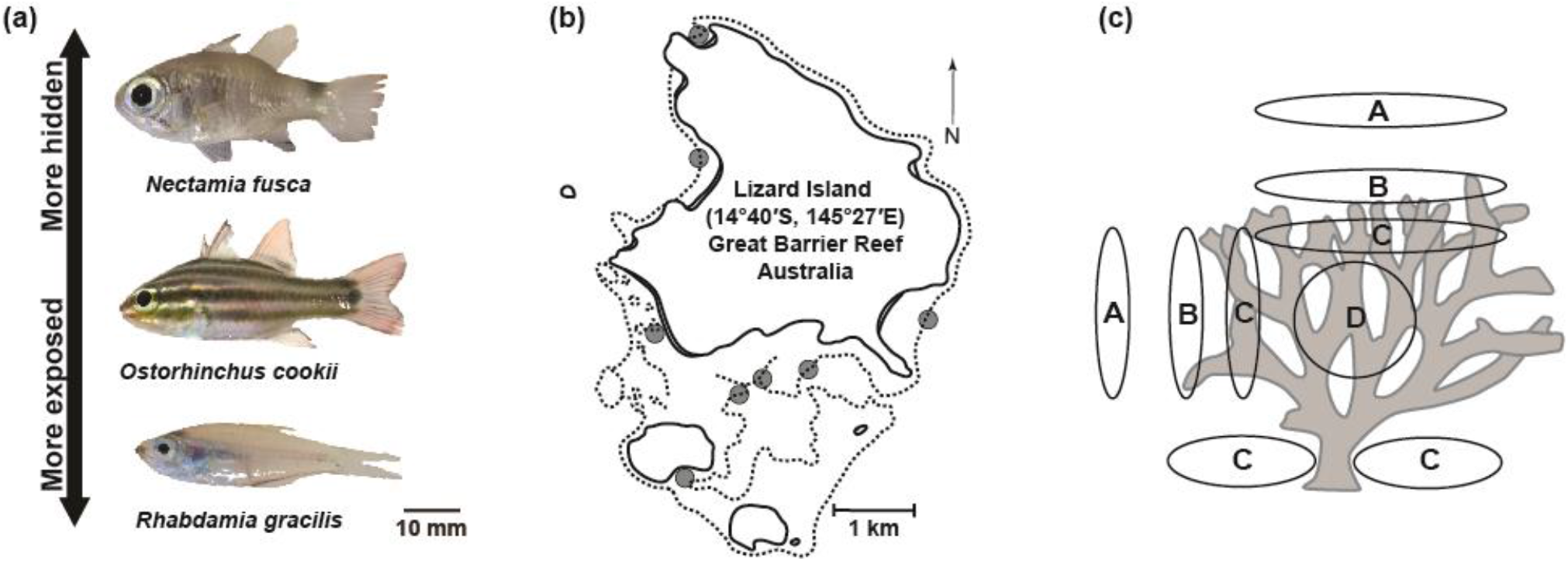
Study species, sampling location and microhabitat assessment. (a) Differences in eye size relative to body length are shown in three cardinalfish species, from top to bottom; Sava Cardinalfish (*Nectamia savayensis*); Cook’s cardinalfish (*Ostorhinchus cookii*); Luminous cardinalfish (*Rhabdamia gracilis*). (b) Overview of the sampling locations (grey circles) around Lizard Island, Queensland, Australia. (c) Schematic of microhabitat classification used for partitioning assessments. Microhabitat A: fully exposed, 1 m above or next to structure (coral head, coral outcrop, boulder); microhabitat B: fully exposed, directly above or adjacent to structure; microhabitat C: semi-hidden (visible from outside), between coral branches, in crevices, under overhangs or ledges; microhabitat D: entirely hidden (not visible from outside), inside coral outcrops, deep inside crevice/cave structures.

**TABLE 1.** Summary of cardinalfish species sampled in microhabitat assessment. n = total individuals counted across all sampling sites and locations. N = number of sampling sites at which each species was found.

| Species | Individuals (n) | Sites (N) |
| --- | --- | --- |
| *Apogon crassiceps* * | 3 | 3 |
| *Apogonichthyoides melas* * | 2 | 2 |
| *Cheilodipterus artus* | 227 | 19 |
| *Cheilodipterus macrodon* | 18 | 7 |
| *Cheilodipterus quinquelineatus* | 1800 | 57 |
| *Fibramia thermalis* | 65 | 3 |
| *Nectamia fusca* | 35 | 3 |
| *Nectamia savayensis* | 50 | 6 |
| *Nectamia viria* * | 10 | 1 |
| *Ostorhinchus compressus* | 104 | 14 |
| *Ostorhinchus cookii* | 44 | 11 |
| *Ostorhinchus cyanosoma* | 2247 | 50 |
| *Ostorhinchus doederleini* | 235 | 24 |
| *Ostorhinchus nigrofasciatus* | 52 | 29 |
| *Ostorhinchus novemfasciatus* * | 3 | 2 |
| *Pristiapogon exostigma* | 44 | 11 |
| *Pristiapogon kallopterus* * | 9 | 3 |
| *Pterapogon cf. mirifica* * | 2 | 2 |
| *Rhabdamia gracilis* | 1410 | 6 |
| *Taeniamia fucata* | 2811 | 13 |
| *Taeniamia zosterophora* | 75 | 3 |
| *Zoramia viridiventer* | 4835 | 15 |
| *Zoramia leptacantha* | 510 | 6 |

\* Omitted from microhabitat partitioning analysis

**FIGURE 2.**
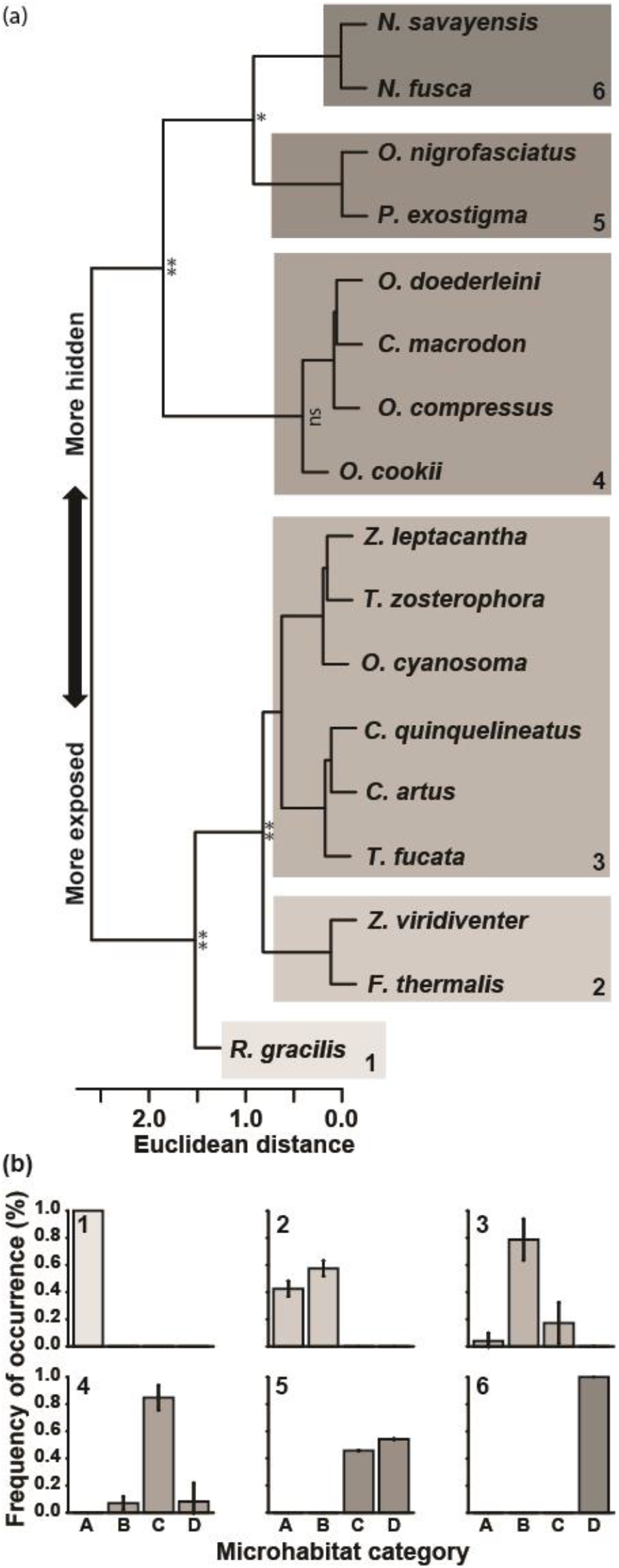
(a) Clustering of microhabitat counts across the four different microhabitat categories (A – D) using a Ward.D2 cluster analysis. Different microhabitat partition clusters are indicated by numbers 1 – 6. Significance levels of bootstrap analysis are designated by: ** = p<0.01, * = p<0.05. (b) Mean (± SD) microhabitat (A – D: A, fully exposed away from structure; B, fully exposed adjacent to structure; C, semi-hidden; D, entirely hidden) distribution of species comprised in each microhabitat partitioning group (1 – 6).

### Relative eye size

Cardinalfish species used for anatomical studies (n = 24) were either collected on the reefs surrounding Lizard Island between February 2015 and April 2017, or obtained through an aquarium supplier (Cairns Marine Pty Ltd, Australia). Fishes from Lizard Island were collected on SCUBA or snorkel using clove oil, hand nets and barrier nets, under the following permits: Great Barrier Reef Marine Park Authority (GBRMPA) Permit G12/35005.1, GBRMPA Limited Impact Permit UQ006/2014 and Queensland General Fisheries Permit 140763. After collection, animals were returned to the lab and anaesthetized using a clove oil solution (10% clove oil, 40% ethanol, 50% sea water) before being euthanized by decapitation.

The standard length (SL) of each individual was measured, eyes were removed from the socket and the horizontal eye diameter was measured to the nearest 0.1 mm using callipers. After removal of the cornea, the lens was extracted and its diameter measured (Table S1). Species were identified based on morphology and colouration and, where possible, subsequently confirmed via RNA-sequencing and by cross-referencing *COI*-sequences to public databases (boldsystems.org) (Luehrmann et al., 2019).

For analyses, relative eye size, lens diameter, eye diameter and standard length were log_10_ transformed. As the ratio between lens diameter and eye diameter was highly proportionate [phylogenetic least squares regression (PGLS), F_1,21_=367.9, r^2^=0.94, p<0.001; Fig. 3a], further comparative analyses were performed on eye diameters only. Relative eye size was calculated as the ratio of log_10_ eye diameter to log_10_ standard length. As data was non-normally distributed (Shapiro-Wilk, w=0.981, p=0.03), analysis of variance for the entire data set was performed using a Kruskal-Wallis test. Differences between cardinalfish relative eye sizes on the genus level were identified through post-hoc pairwise comparisons [Dunn-test; (Dunn, 1961)]. To account for multiple comparisons, p-values were adjusted using a Bonferroni corrections (Table S2). Genera for which three or fewer individuals were measured were omitted from the analysis (Table S1).

**FIGURE 3.**
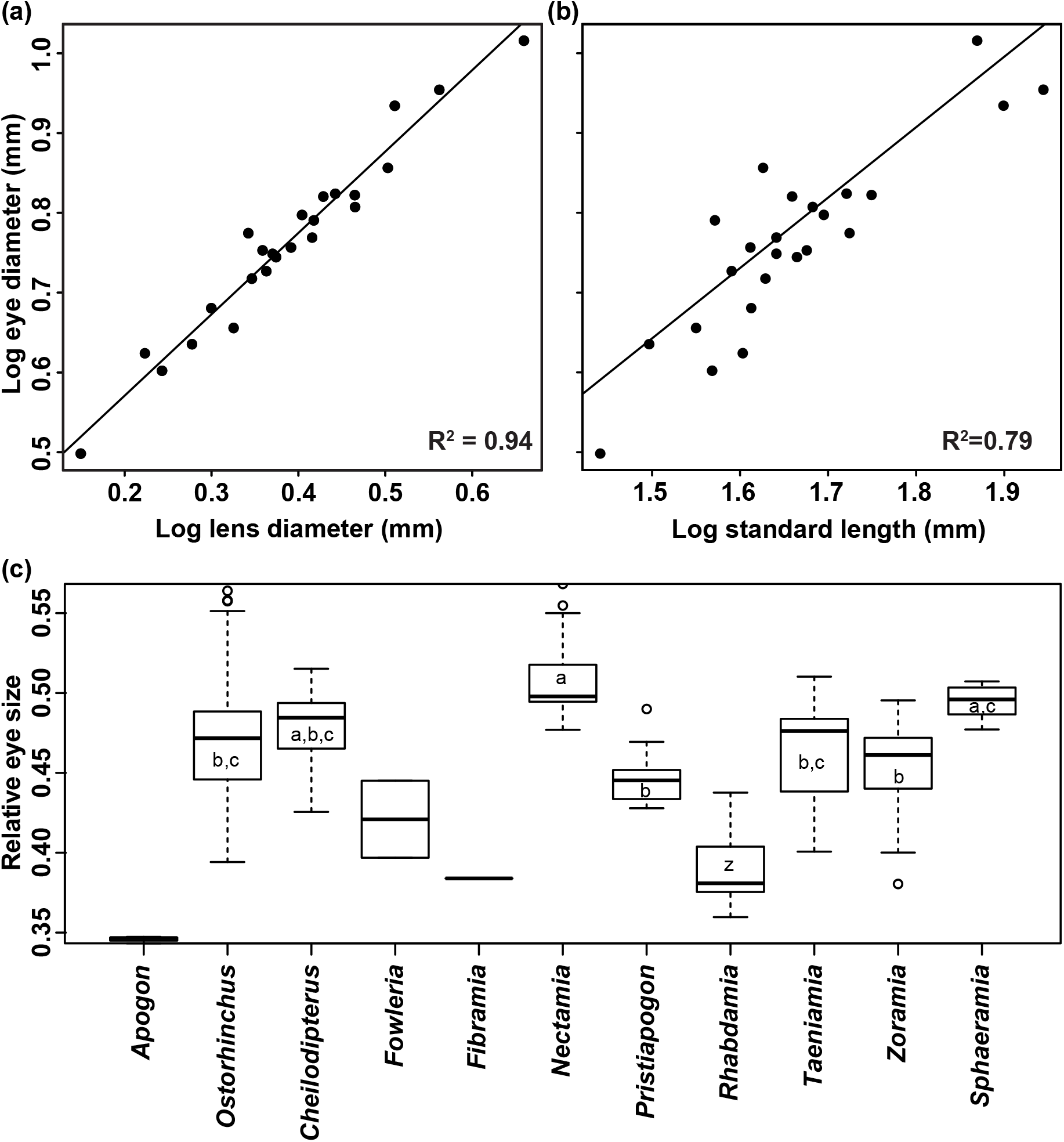
Relationships of (a) horizontal eye diameter and lens diameter, and (b) eye diameter and standard length. Fitted lines represent the phylogenetically corrected linear regressions using PGLS. (c) Comparison of relative eye size by genus as per Mabuchi et al. (2014). Different letters indicate significant differences based on Dunn’s post-hoc tests. Genera without letters were excluded from the analysis (see Table S1).

### Retinal cell topography

In five cardinalfish species, enucleated eyecups were fixed in 4% paraformaldehyde (PFA) in 0.1 M Phosphate Buffered Saline (PBS) at room temperature for at least 24 hours, then stored at 4°C. Retinal wholemounts were prepared following the methods outlined in Coimbra et al. (2006) and Ullmann et al. (2012). Few individuals were investigated in this study due to the challenge in processing such small eyes using this method. However, previous studies (de Busserolles et al., 2014a,b) have shown a low intraspecific variability in retinal topography in fishes, and the analysis of two individuals of *Ostorhinchus doederleini* (Fig. S2), suggests low intraspecific variability in cardinalfishes also. Using the stereological software StereoInvestigator (Microbrightfield), topographic distribution of photoreceptors and ganglion cells was assessed using the optical fractionator technique designed by West et al. (1991) and modified by Coimbra et al. (2012). The counting frame and grid sizes were carefully chosen to maintain the highest level of sampling and achieve an acceptable Schaeffer’s coefficient of error (CE <0.1; Glaser and Wilson, 1998; Slomianka and West, 2005) following the sampling protocols described in de Busserolles et al. (2014a,b) (see Table S3 for a summary of counting parameters). For ganglion cell analysis, displaced amacrine cells were included in the counts as they were difficult to distinguish from ganglion cells based on morphological criteria alone. The inclusion of amacrine cells in the analysis has previously been shown not to influence the overall topography of fish retinae (e.g. Collin and Pettigrew, 1988c). Topographic maps were constructed using R v3.1.2 (R Core team, 2014) with the results exported from StereoInvestigator according to Garza-Gisholt et al. (2014). The upper limit spatial resolving power (SRP), expressed in cycles per degree (cpd), was estimated for each individual using the ganglion cell peak density as described by Collin and Pettigrew (1989). Note that, since amacrine cells were included in the ganglion cell counts, SRPs will be slightly overestimated.

### Opsin gene expression, activity patterns and foraging mode

We used proportional opsin gene expression data from our previous work on 26 cardinalfish species collected from the same locations (Luehrmann et al., 2019, Table S5). These included all of the species used for microhabitat partitioning analysis (Table 1) and those for which relative eye sizes and retinal topography maps were obtained (Tables S1, S3). We also characterised each species as being nocturnally or diurnally active, and their foraging mode as exclusively benthivorous, benthivorous and planktivorous, or exclusively planktivorous based on previously published research (see Table S5 for references).

### Phylogenetic comparative analyses

We tested whether the ecological parameters (microhabitat use, feeding mode, activity period) correlated with visual system design (relative eye size, proportional opsin gene expression) of cardinalfishes using PGLS. For comparative analyses, we used the cardinalfish phylogeny from Luehrmann et al. (2019). Each predictor was independently tested against each dependent variable and no correlations between predictors were assessed due to different sample sizes. To account for multiple testing, p-values were adjusted using Bonferroni corrections: for eight tests each in the cases of proportional opsin gene expression versus microhabitat and feeding mode, respectively; and for seven tests each in the cases of proportional opsin gene expression versus activity period and versus relative eye size, respectively (Fig. 5, Tables S5, S6). Analyses were performed in R version 3.1.2 (R Core team, 2014) using the CAPER package (Orme, 2013).

## Results

### Microhabitat distribution

We found marked variability in abundance and microhabitat distribution among different cardinalfish species (Table 1; Fig. S1). Several species displayed microhabitat specialisations (e.g., *Rhabdamia gracilis*, *Nectamia savayensis*, *N. fusca*, *Taeniamia zosterophora*, *Zoramia leptacantha*, *Ostorhinchus compressus*), while others showed more generalist microhabitat preferences (e.g., most *Cheilodipterus* and *Ostorhinchus* species) (Fig. S1).

Hierarchical cluster analysis revealed that cardinalfishes can be broadly classified into six habitat specialisation clusters (Fig. 2a), based on the microhabitat(s) they were most frequently found in (Fig. 2b). Cluster 1 contained only *R. gracilis*, which was exclusively found away from, but within 1-2 m of, the reef structure in midwater (microhabitat A; bootstrap, p<0.01). Cluster 2 contained species (*Fibramia thermalis*, *Zoramia viridiventer*) found mostly exposed, but located close to structure, e.g. hovering above the tips of branching corals (microhabitat A or B, p<0.01). Cluster 3 consisted of species of the genera *Taeniamia*, *Ostorhinchus*, *Cheilodipterus* and *Zoramia* that were predominantly exposed, but were sometimes found in cover (microhabitat B, sometimes A or C, p<0.01). Cluster 4 consisted of species (*Ostorhinchus cookii*, *O. compressus* and *O. doederleini*) that were found predominantly in cover, either at the bottom of corals underneath branches, beneath rock ledges, or between the tips of branching corals (microhabitat C, p<0.01). They were, however, easily spotted from outside. Cluster 5 comprised species (*Pristiapogon exostigma*, *O. nigrofasciatus*) that were always hidden, e.g. under ledges, between coral branches, or inside caves, where they were sometimes hard to spot (microhabitat C or D, p<0.05). Finally, cluster 6 comprised species found exclusively hidden inside the reef matrix, mostly deep inside branching corals (*Nectamia savayensis*, *N. fusca*; microhabitat D, p<0.05).

### Relative eye size

Eye diameter was proportional to body size (PGLS, F_1,21_=85.11, r^2^=0.79, p<0.001), but showed considerable variation between species (Kruskal-Wallis, χ^2^=116.434, df=10, p<0.001; Fig. 3b, see Table S1 for an overview of morphometric measurements). Post-hoc pairwise comparisons of relative eye size at the genus level furthermore revealed three distinct size categories in this family (Fig. 3c, Table S2). Members of the genus *Nectamia* had the largest eyes relative to their body sizes. Species of the genera *Ostorhinchus*, *Cheilodipterus*, *Pristiapogon*, *Taeniamia* and *Zoramia*, on the other hand, had intermediate sized eyes, while showing greater variability. *Ostorhinchus* species, in particular, showed a wide range of eye-diameter-to-standard-length ratios. *Sphaeramia nematoptera* had consistently large eyes, but not statistically larger than *Ostorhinchus* (Dunn, z=−2.121, p=0.036), *Cheilodipterus* (z=−1.04, p=0.2) and *Taeniamia* genera (z=2.103, p=0.035) due to the broad range of eye sizes in these groups. *R. gracilis* had the smallest eyes overall, even when compared to the *Zoramia* (z=−4.373, p<0.001) and *Taeniamia* (z=−4.628, p<0.001) species, which had the second smallest relative eye sizes. *Apogon crassiceps* appeared to have even smaller eyes, however, as only three specimens were sampled, this species was omitted from the analysis. In summary, species that have intermediate sized eyes showed greater variability than species with consistently large or consistently small eyes (Fig. 3, Tables S1, S5).

Microhabitat partitioning correlated with relative eye size (PGLS, F_5,11_=7.66, p=0.02; Fig. 5d, Table S6). Relative eye size showed a positive correlation to decreased microhabitat exposure, with *R. gracilis* (microhabitat A) having the smallest, and *N. savayensis* and *N. fusca* (microhabitat D) having the largest eyes. Interestingly, species occurring predominantly in both completely hidden (microhabitat D) and partially hidden (microhabitat C) microhabitats (cluster 5), had surprisingly small eyes (*P. exostigma*, *O. nigrofasciatus*; Fig. 5d). Relative eye size showed no significant correlation to either activity period or feeding mode (Table S6).

### Retinal neural cell topography

Retinal cell topographies differed between genera, and in one case between species of the same genus (*Ostorhinchus notatus* differed from *O. cyanosoma* and *O. doederleini*; Fig. 4). Topographic maps of ganglion cell densities revealed two specialization types, one characterized by increased cell density in the central and temporo-ventral part of the retina (*R. gracilis*, *T. fucata*), and one characterized by increased cell density in the central part of the retina (area centralis), which extends into a weak horizontal streak (*O. cyanosoma*) (Fig. 4).

**FIGURE 4.**
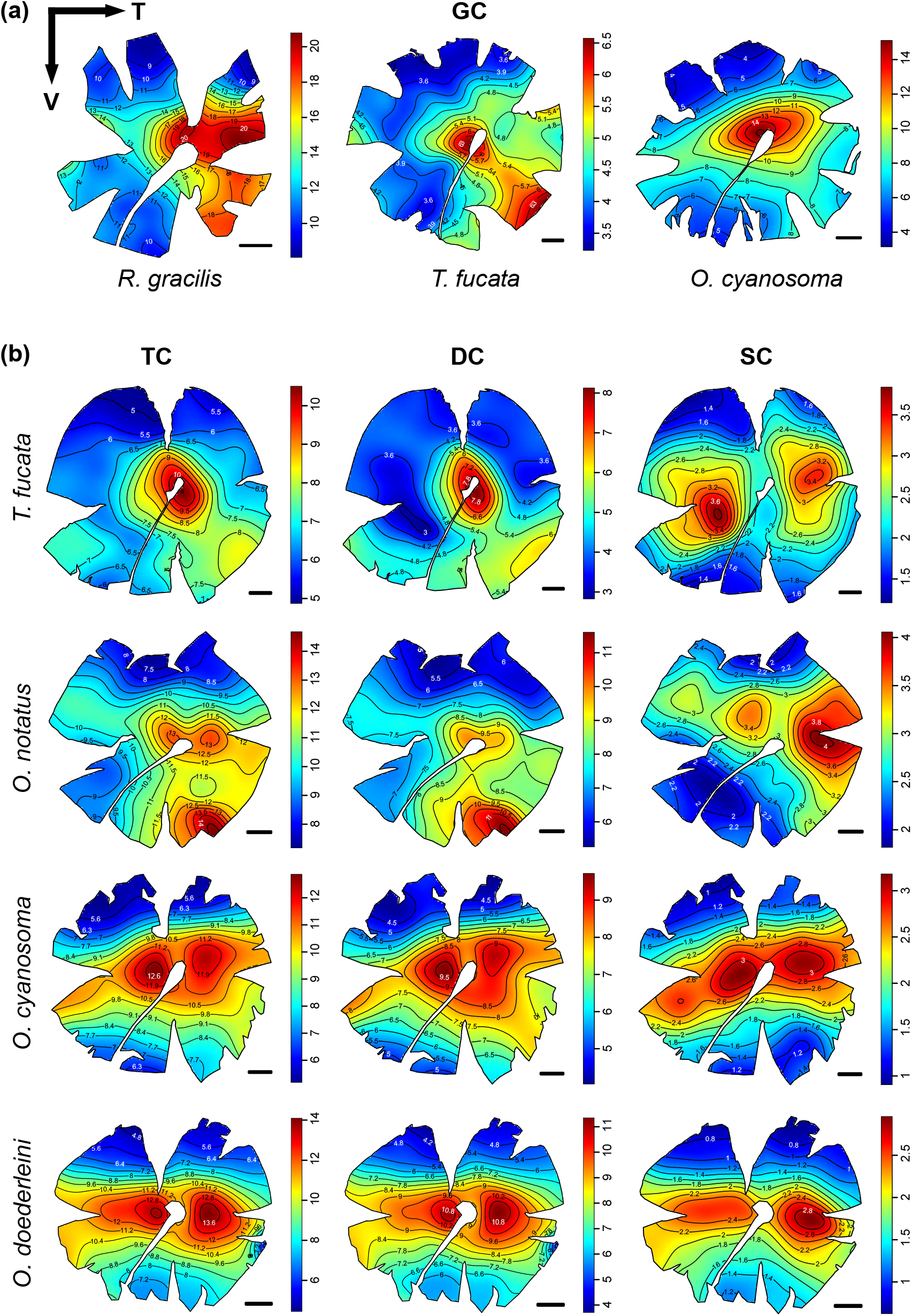
Topographic distribution of retinal neural cells in different cardinalfish species. (a) Ganglion cells (GC), (b) photoreceptors (total cone TC, double cone DC, and single cone SC). For two species, both ganglion and photoreceptors cells have been mapped. Black lines represent isodensity contours and values are expressed in densities × 10^3^ cells/mm^2^. Arrows indicate the orientation of the retinas: V = ventral, T = temporal. Scale bars = 1 mm.

Photoreceptors were mostly arranged in a square mosaic pattern composed of one single cone surrounded by four double cones. However, in some species, this pattern was not consistent over the entire retina with some areas showing more irregular single cone patterns (Fig. S4). Topographic maps of total cone and double cone densities were nearly identical and revealed three specialisation types (Fig. 4): an area centralis (*T. fucata*); an increase in cell density in a large area of the central and temporo-ventral part of the retina and extending into a weak horizontal streak (*O. notatus*); and a pronounced horizontal streak with two areae centralis (*O. cyanosoma* and *O. doederleini*). For the species for which both ganglion cells and photoreceptors were investigated (*T. fucata*, *O. cyanosoma*), total photoreceptor cell distributions were similar to the ganglion cell distributions. Single cone topography was noticeably different from double cone topography in two species (*T. fucata*, *O. notatus*). These two species showed the highest proportion of single cones of all species investigated and *T. fucata* had irregular single cone patterns (Fig. S4). In *T. fucata*, single cone distribution showed two large areas of increased cell density in the nasal and temporal part of the retina, while double cone distribution showed a single area centralis around the optic nerve (Fig. 4). In *O. notatus*, single cone density was highest in the temporal region of the retina, extending into a horizontal streak, whereas double cone density was highest in the temporo-ventral part of the retina (Fig. 4).

No clear relationship between retinal topography and microhabitat use and/or activity pattern could be identified. Species occupying the same microhabitat partition had very different topographies (e.g. *T. fucata* and *O. cyanosoma*; Fig 4, Table S5) and species mainly differing in activity pattern had similar topographies (e.g. *O. cyanosoma* and *O. doederleini*; Fig 4, Table S5). However, topography and feeding mode seemed to correlate, with pure planktivores having an area temporo-centralis (*R. gracilis*, *T. fucata*), while generalists (i.e., benthic and pelagic feeders) possessed streaks (*O. cyanosoma*, *O. doederleini*; Fig. 4, Table S5). Moreover, the type of specialisation appeared to follow the phylogeny, with closely related species having similar retinal topographies (Fig. S3).

Total cell numbers and densities of both ganglion cells and photoreceptors varied between species, but appeared to be of similar order of magnitude (Table S4). Peak ganglion cell density ranged from 8,289 cells/mm^2^ in *T. fucata* to 23,051 cells/mm^2^ in *R. gracilis*. However, spatial resolving power was similar in the three species assessed, with an average of 7.6 cycles per degrees.

### Opsin gene expression

We found that *SWS2B* (PGLS, F_5,11_=9.283, p=0.009), *RH2A* (F_5,11_=10.31, p=0.006) and *LWS* (F_5,11_=11.17, p=0.004) cone opsin expression, and the rod-opsin to cone-opsin ratio (F_5,11_=20.37, p<0.001) correlated with microhabitat partitioning (Fig. 5c, Tables S5, S6). *LWS* was highly expressed in *Nectamia* species, and, at lower levels, in *Fibramia*, but in virtually no other species. Therefore, *LWS* expression appeared to be high in species exclusively occupying hidden microhabitats (microhabitat D). *SWS2B*, in contrast, was highly expressed in species exclusively occupying exposed microhabitats (microhabitat A, *R. gracilis*; microhabitat B, *Z. viridiventer*). *RH2A*, while expressed in all species, showed the highest expression (> 80% of double cone gene expression) in species occupying in-between microhabitats (clusters 2 – 5). *RH1* expression correlated negatively with microhabitat exposure, and thus was lowest in *R. gracilis*, followed by *F. thermalis* and *Z. viridiventer*. However, in all remaining species, *RH1* expression was nearly identical and accounted for > 90% of total opsin expression (Fig. 5c, Table S5). Activity period, feeding mode and relative eye size were not significantly related to any opsin expression profiles tested (Table S6).

**FIGURE 5.**
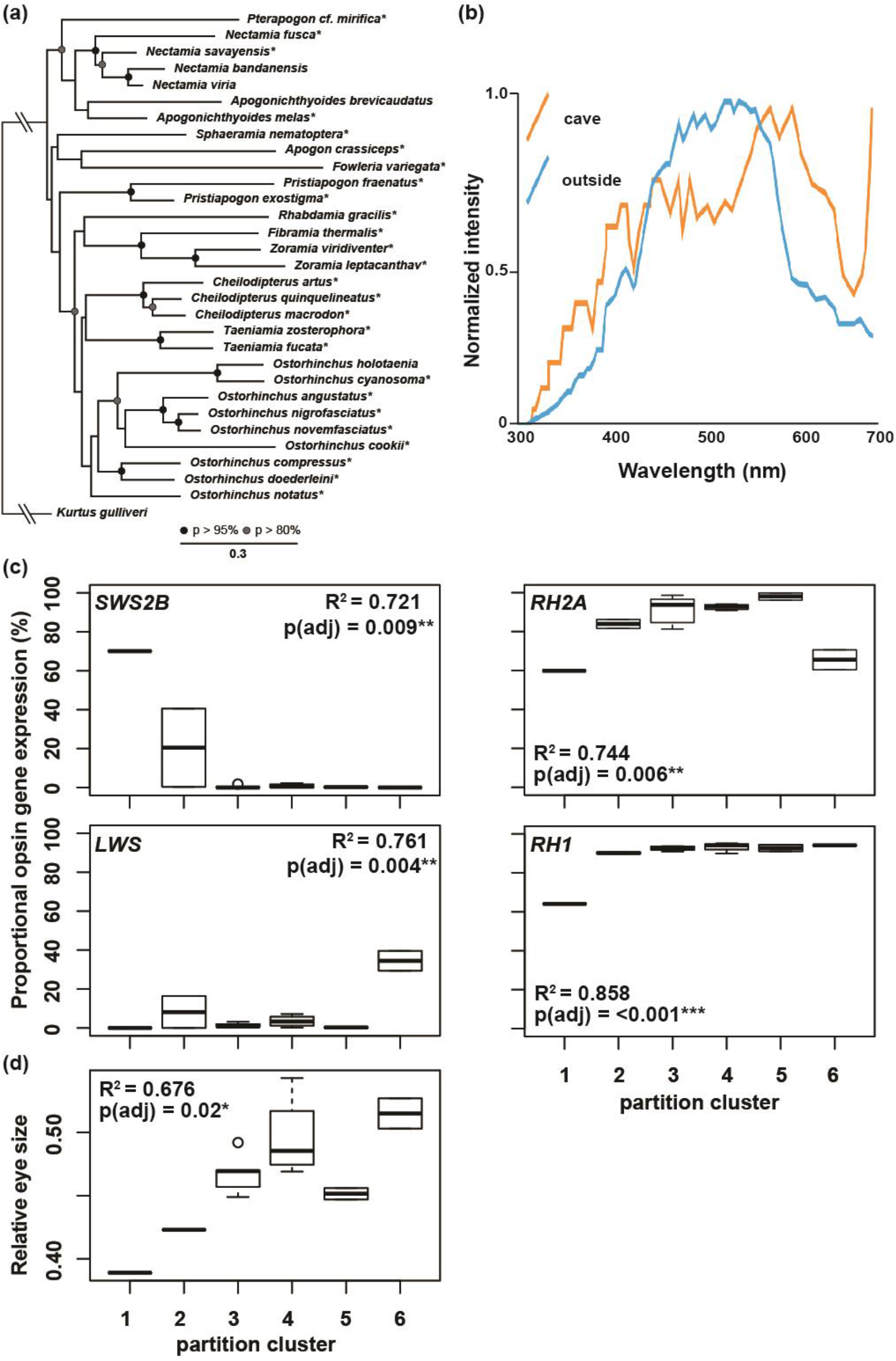
Phylogenetic comparative analysis (PGLS) of cardinalfish visual system characteristics in relation to ecological parameters and visual system specialisations. (a) Cardinalfish phylogeny used for PGLS analysis (see Luehrmann et al., 2019). Asterisks indicate species used and spheres indicate maximum likelihood support values. (b) Light environment in different microhabitats on coral reefs at Kaneohe Bay, Hawaii (after Marshall et al., 2003). Orange line, light inside a cave 1 m recessed. Blue line, light on the reef outside the cave. Measurements in relative photons/sr/nm. (c) Proportional opsin gene expression and (d) relative eye size in cardinalfishes categorised by microhabitat partitioning clusters. Bonferroni adjusted p-values are shown where: * < 0.05, ** < 0.01, *** < 0.001.

## Discussion

Our results show that microhabitat partitioning among different cardinalfish species correlates with adaptations in their visual systems. Specifically, we found that a reduction in relative eye size, and therefore light sensitivity, was present in species from exposed microhabitats compared to species found hidden in the substrate. Opsin gene expression was also related to microhabitat use, with exposed species expressing a shorter-wavelength-shifted and hidden species expressing a longer-wavelength-shifted cone opsin palette, probably reflecting each microhabitat’s light colour (Fig. 5b). As to whether microhabitat partitioning could also explain differences in retinal neural cell topography, our results were inconclusive. Instead, a possible link between retinal topography and feeding mode was identified.

However, these trends were driven mainly by the few species showing extreme forms of adaptations to light conditions in microhabitats situated at the extreme ends of the light intensity and colour spectrum (microhabitats A and D). Most other species fall somewhere in between, with visual systems that seem suited to a broader colour and intensity range (see Figure 2). Since selection pressure is expected to be relaxed under those conditions, phylogenetic inertia may play a major role in shaping the visual systems in in-between cardinalfishes. This is also supported by the strong phylogenetic signal (Pagel’s λ) when correlating relative eye size, *SWS2B* expression and *LWS* expression with microhabitat (Table S6).

Microhabitat partitioning behaviour occurs in many animals due to resource competition, such as for food, suitable mating sites, or shelter from predators (e.g., Ross, 1986). With few exceptions, most cardinalfish species forage nocturnally and away from their diurnal refuge sites (Barnett et al., 2006; Marnane and Bellwood, 2002). For these species, partitioning at their diurnal refuge sites is unlikely to be due to competition for food. An exception can be found in species with increased expression of violet opsin (*SWS2B*) that also occur in exposed microhabitats and feed diurnally (*R. gracilis*). They may benefit from shorter-wavelength-shifted visual systems compared to other species (Luehrmann et al., 2019) in feeding contexts in midwater microhabitats as UV-sensitivity can aid planktivory (e.g. Flamarique, 2016). Moreover, *R. gracilis* showed decreased expression of rod opsin (*RH1*), which supports its diurnal lifestyle. With the exception of *R. gracilis*, cardinalfishes are well adapted to dim-light by means of increased *RH1* expression compared to diurnal reef fishes (Luehrmann et al., 2019). However, even among those nocturnally foraging cardinalfishes the repertoire of cone opsins they use is on par with those of diurnal coral reef fishes (Luehrmann et al., 2019). It remains to be tested whether cardinalfishes are capable of dim-light colour vision which could improve night time foraging efficiency, as reported from some gecko and anuran species (Kelber and Roth, 2006).

Cardinalfishes are heavily preyed on, making efficient defence mechanisms critical for their survival (e.g., Beukers-Stewart and Jones, 2004). Consequently, competition for shelter may drive microhabitat partitioning in this family, with those forced into the open needing to develop other means of protection. Indeed, several species generally found in microhabitats away from structure (microhabitat A and B), such as *R. gracilis*, *Z. viridiventer*, *T. zosterophora*, or *Z. leptacantha*, are of silvery-translucent or pale appearance, providing excellent camouflage when viewed against a blue water background (Marshall and Johnsen, 2011; Marshall et al., 2018). These species also form large schools, possibly further reducing predation risk (Pitcher, 1986). In contrast, species found in more sheltered microhabitats, e.g., *O. doederleini*, *O. cookii*, *O. compressus*, or those always hidden inside the reef matrix, e.g., *N. savayensis*, are darker in overall body colour, or like most *Ostorhinchus* species, have dark horizontal stripes. These species are also often solitary or live in smaller groups (Randall et al., 1990). Light inside caves and crevices on tropical coral reefs is dim and red-shifted, the latter presumably caused by encrusting red-algae or other red-pigmented encrusting organisms, like sponges (Fig. 5b, Marshall et al., 2003). Increased expression of *LWS* (red-sensitive) opsin, along with a complete absence of *SWS2B* expression, particularly in *Nectamia* species, may be an adaptation to the dim and/or red-shifted light conditions present in these microhabitats.

A longer-term effect of visual adaptations to different microhabitat light spectra may be enhanced con- or heterospecific recognition in context of relevant behaviours under the given light environments, such as mate choice and/or predator avoidance, as these are carried out during the day (Kuwamura, 1983; Kuwamura, 1985; Saravanan et al., 2013). Mate colouration and visual adaptation to it may be important for sexual selection, as seen in cichlid speciation (Terai et al., 2006; Seehausen et al., 2008). Colour or pattern based heterospecific recognition, on the other hand, may be essential for the assortative aggregation behaviour of cardinalfishes (Gardiner and Jones, 2005; Greenfield and Johnson, 1990), or predator recognition. Indeed, most cardinalfish species are colourful and many sport stripe patterns, as well as violet and/or UV-markings, which are visible through their relatively UV-transparent ocular media and SWS photopigments (Marshall, 2000; Siebeck and Marshall, 2001). However, unlike some reef fish (e.g., damselfishes) in which UV facial patterning can be slightly different between males and females and even between individuals (Siebeck et al., 2010), cardinalfishes do not exhibit sexual dimorphism: males can only be distinguished from females by their enlarged jaws during egg incubation in breeding season (Barnett and Bellwood, 2005).

The relationship of eye sizes to habitat exposure and brightness found here, with species occupying the most sheltered and dimmest microhabitats having the largest eyes and species occupying the least sheltered and brightest microhabitats having the smallest eyes, is consistent with previous studies in fishes (Schmitz and Wainwright, 2011), and confirm the importance of light availability as a driver of eye size evolution. However, some species (*P. exostigma*, *O. nigrofasciatus*) found in both completely (microhabitat D) and partially hidden microhabitats (microhabitat C) had surprisingly small eyes, suggesting that other factors, such as phylogeny (Table S6, de Busserolles et al., 2013) or predation (Beston et al., 2017), may have influenced their eye size evolution.

In fishes, the retinal topography usually reflects their habitat type, with visual streaks found in species living in open environments with an uninterrupted view of the horizon, and areae centralis found in species living in more enclosed environments (Terrain theory by Collin and Pettigrew, 1988a; Collin and Pettigrew, 1988b; Hughes, 1977). Here, the lack of a relationship between microhabitat type and retinal topography in cardinalfishes could be explained by several factors: 1) we focused on the cone topography, but many of the species studied here are nocturnal and therefore may rely more on their rod photoreceptors; 2) habitat partitioning in this study was only assessed during the day and no data is available on habitat partitioning at night. Hence, it is possible that a correlation between microhabitat partitioning and retinal topography exists, but that it was missed due to different activity periods of species; 3) retinal topography in cardinalfishes may carry a strong phylogenetic signal (Fig S3). A phylogenetic constraint in retinal topography was first suggested in mammals (Stone 1983) and has since been observed in a number of animals, including in deep-sea fishes (de Busserolles et al., 2014a); 4) instead of diurnal microhabitats, retinal topography in cardinalfishes may be influenced by their feeding ecology. The streak in benthivorous/plantivorous species may allow them to scan a broad area of the sand-water interface, providing high acuity while minimising eye movements (Collin and Pettigrew, 1988b). On the other hand, a higher density of cells in the temporo-ventral part of the retina in obligate planktivores may provide higher acuity to distinguish prey situated in front and above in the water column (Collin and Pettigrew, 1988a; de Busserolles et al., 2014a).

## Conclusion

Our findings suggest that microhabitat partitioning is a contributing factor to visual system diversification in cardinalfishes, specifically those adapted to extreme microhabitats, whereas many show visual systems suited to broader microhabitat conditions. The colour vision of these nocturnal foragers is presumably linked to daytime activities in and around coral heads and the close-set nature of these social and other activities may in particular drive this site-system co-adaptation. While there remains much to learn around the colours and their use in these engaging fish, our findings indicate that the availability of diverse microhabitats contributes to evolutionary sensory diversification among diverse ecosystems, such as tropical coral reefs, and perhaps in similar ways in terrestrial systems.

## Authors’ contributions

ML, FC, KLC, FdB, NJM conceived the ideas and designed the methodology; ML and FC collected the data; ML analysed the data; ML led the writing of the manuscript; all authors contributed critically to the drafts and gave final approval for publication.

## Acknowledgements

We thank the staff at Lizard Island Research Station for support during field work and Cairns Marine Pty Ltd for sourcing and supplying additional fish. This work was supported by an Australian Research Council Discovery Project (DP150102710) awarded to KLC and JM, an ARC Laureate Fellowship (FL140100197) to JM, a UQ Development Fellowship to FC, and FdB was funded by an ARC DECRA (DE180100949).

## Data Accessibility

All data is given either in the main manuscript, the supplementary material or is available through the publications cited accordingly.

