## Supplementary information for "Microhabitat partitioning correlates with opsin gene expression in coral reef cardinalfishes (Apogonidae)"

**FIGURE S1.** Proportion of individuals per species counted in each defined microhabitat category across sites. N = number of sampling sites at which each species was found. n = total number of individuals counted. Also see Figure 1 for microhabitat categorisation.

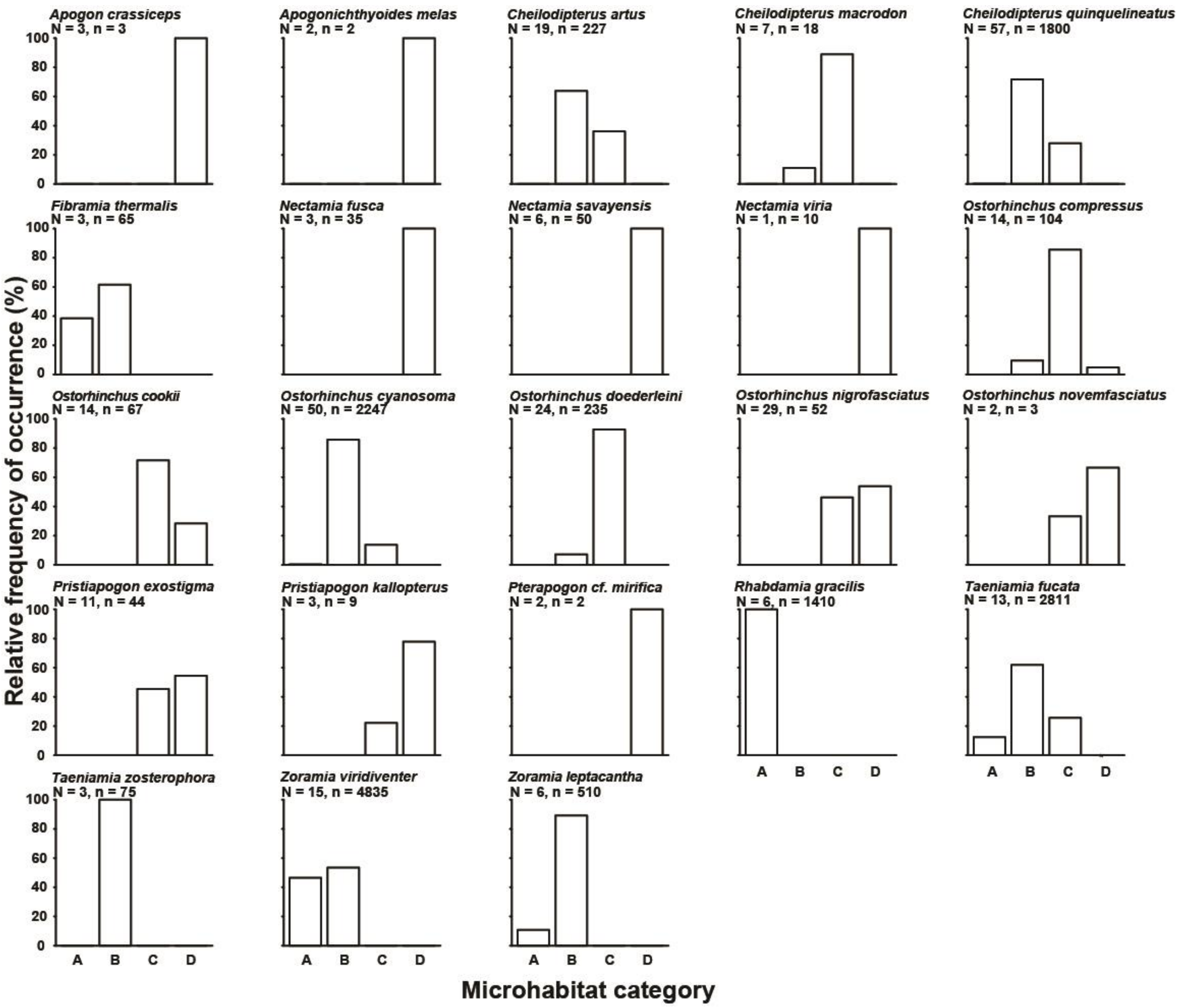

**FIGURE S2.** Topographic distribution of photoreceptor cells (total cones TC, double cones DC, single cones SC) in two individuals of *Ostorhinchus doederleini* highlighting a low intraspecific variability. Black lines represent isodensity contours and values are expressed in densities x $10^3$ cells/mm$^2$. The arrows indicate the orientation of the retinas, V = ventral, T = temporal. Scale bars = 1 mm.

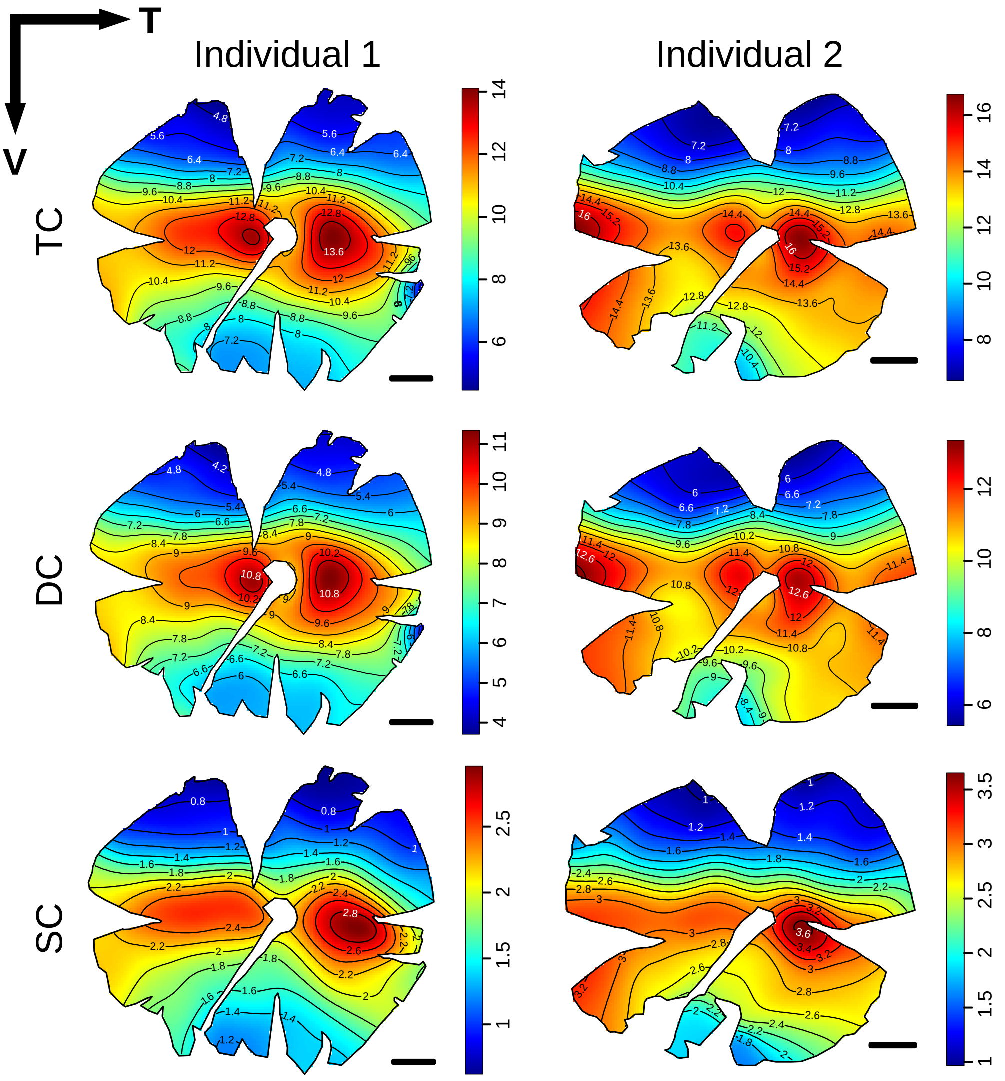

**FIGURE S3.** Type of retinal specialisation in the five cardinalfish species analysed in this study plotted against the cladogram of the family. Cladogram made from the phylogeny presented in Figure 5a. A = area, S = streak.

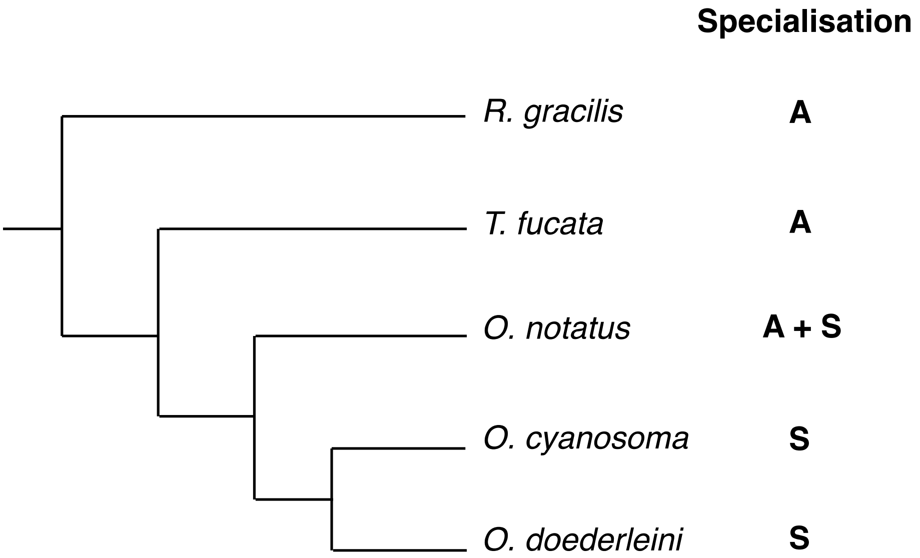

**FIGURE S4.** Retinal photoreceptor mosaic patterns found in cardinalfishes. (a) square pattern, (b) chaotic pattern (after Collin and Shand 2003). (c) Light microscopy view of the central-temporal part of the retina in *O. doederleini*. (d) Light microscopy view of the nasal part of the retina in *T. fucata*. (e) and (f) Location of photoreceptor mosaic patterns shown superimposed over total photoreceptor distribution maps from Fig. 4 (main manuscript). S = square pattern, C = chaotic pattern.  Black lines represent isodensity contours. Values are expressed in densities x $10^3$ cells/$mm^2$. The arrows indicate the orientation of the retinas, V = ventral, T = temporal. Scale bars = 1 mm.

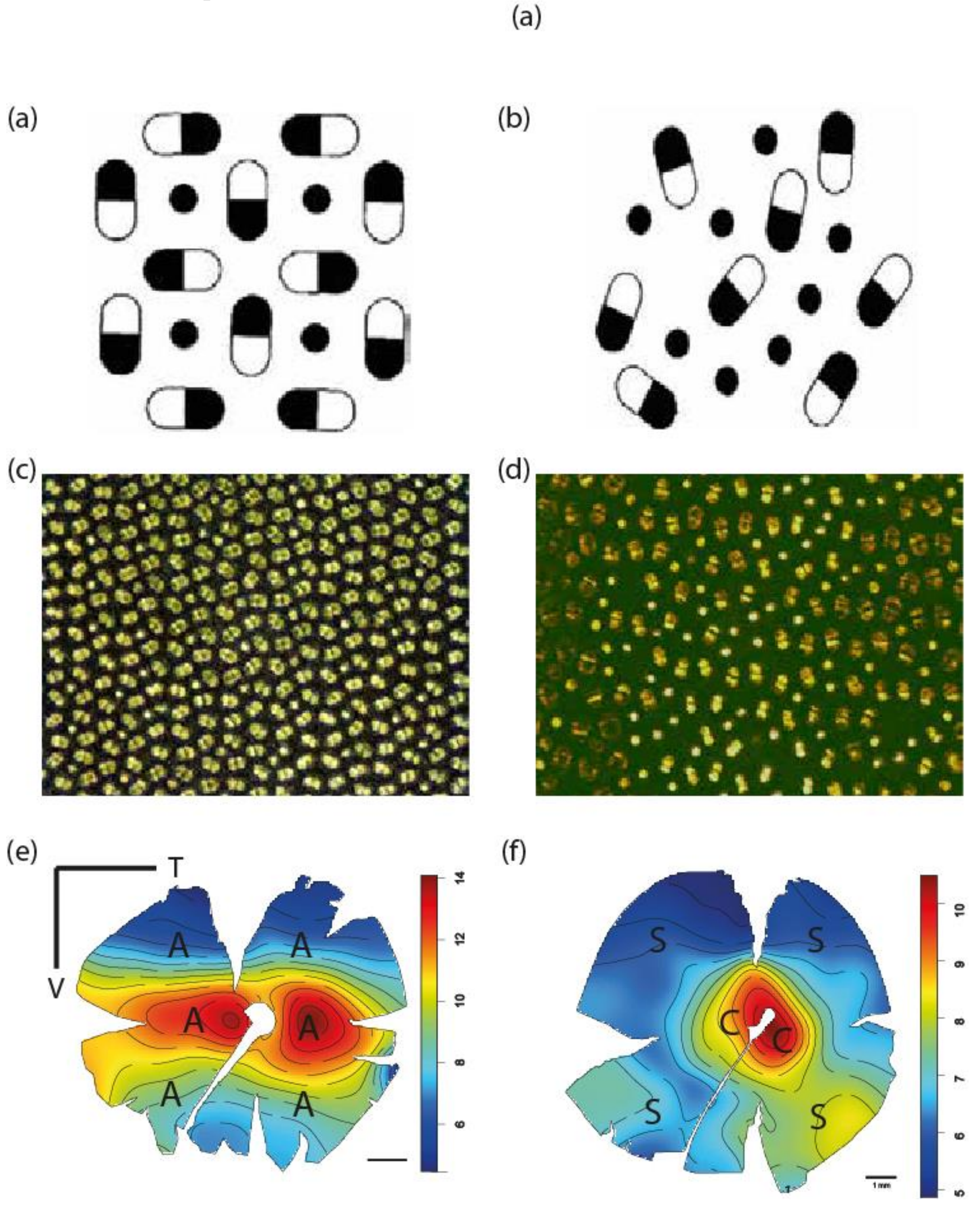

**TABLE S1.** Range of standard length (SL) and horizontal eye diameter measured in sampled cardinalfishes.

| **Species** | | **n** | **SL (mm)** | **Eye diameter (mm)** | **Lens diameter (mm)** |
|---|---|---|---|---|---|
| *Apogon* | *crassiceps* | 3 | 26 - 30 | 3.1 - 3.3 | 1.25 - 1.55 |
| *Cheilodipterus** | *artus* | 4 | 66 - 96 | 6.95 - 10.5 | 2.95 - 3.40 |
| | *macrodon* | 1 | 88 | 9 | 3.65 |
| | *quinquelineatus* | 6 | 46 - 69 | 5.1 - 8.6 | 2.5 - 3.25 |
| *Fibramia* | *thermalis* | 1 | 37 | 4 | 1.75 |
| *Fowleria* | *variegata* | 2 | 35 - 48 | 4.1 - 5.6 | 1.9 - 2.2 |
| *Nectamia** | *fusca* | 6 | 35 - 58 | 6.1 - 10.1 | 2.6 - 4.2 |
| | *savayensis* | 22 | 26 - 53 | 4.8 - 9.1 | 2.0 - 4.3 |
| | *compressus* | 11 | 63 - 85 | 8.8 - 12.3 | 3.7 - 5.3 |
| *Ostorhinchus** | *cookii* | 21 | 28 - 63 | 4.3 - 8.1 | 3.95 - 1.8 |
| | *cyanosoma* | 19 | 31 - 48 | 4.2 - 6.7 | 1.9 - 3.0 |
| | *doederleini* | 15 | 33 - 52 | 4.9 - 6.4 | 2.0 - 2.95 |
| | *holotaenia* | 3 | 26 - 41 | 3.9 – 5.1 | 1.8 – 2.1 |
| | *nigrofasciatus* | 14 | 30 - 57 | 4.25 - 6.75 | 1.75 - 2.95 |
| | *notatus* | 3 | 50 - 55 | 6.5 - 6.9 | 2.8 |
| | *novemfasciatus* | 9 | 35 - 49 | 4.15 - 6.6 | 1.85 - 2.65 |
| *Pristiapogon** | *exostigma* | 12 | 40 - 77 | 5.1 - 8.4 | 2.0 - 3.75 |
| | *fraenatus* | 1 | 53 | 5.95 | 2.2 |
| *Rhabdamia** | *gracilis* | 24 | 34 – 47 | 3.6 – 5.3 | 1.35 – 1.9 |
| *Sphaeramia** | *nematoptera* | 7 | 41 - 49 | 5.95 - 7.2 | 2.35 - 3.1 |
| *Taeniamia** | *fucata* | 5 | 37 - 60 | 4.25 - 7.85 | 1.6 - 3.6 |
| | *zosterophora* | 12 | 40 - 56 | 5.05 - 7.8 | 2.05 - 3.2 |
| *Zoramia** | *leptacanthas* | 22 | 33 - 49 | 4.6 - 6.9 | 1.8 - 2.96 |
| | *viridiventer* | 11 | 31 - 44 | 3.95 - 5.5 | 1.67 - 2.5 |

n = individuals measured

* Used to identify relative eye size differences on the genus level (see also Fig. 4c, Table S2)

**TABLE S1**. Results of post-hoc pairwise comparisons of relative eye size in the investigated cardinalfish genera using Dunn's post-hoc test and p-value adjustment according to Hochberg (1995). Significant p-values are shown in bold.

| **Genus** | | *Cheilodipterus* | *Nect.* | *Osto.* | *Pristi.* | *Rhabd.* | *Sphaer.* | *Taeni.* |
|---|---|---|---|---|---|---|---|---|
| *Nectamia* | z | -1.912 | | | | | | |
| | p | 0.051 | | | | | | |
| *Ostorhinchus* | z | 1.03 | **4.688** | | | | | |
| | p | 0.199 | **0** | | | | | |
| *Pristiapogon* | z | **2.427** | **4.989** | 2.253 | | | | |
| | p | **0.019** | **0** | 0.027 | | | | |
| *Rhabdamia* | z | **5.242** | **9.307** | **6.92** | **2.655** | | | |
| | p | **0** | **0** | **0** | **0.012** | | | |
| *Sphaeramia* | z | -1.04 | 0.419 | -2.121 | **-3.193** | **-5.614** | | |
| | p | 0.2 | 0.357 | 0.036 | **0.0028** | **0** | | |
| *Taeniamia* | z | 1.141 | **3.649** | 0.432 | -1.5 | **-4.628** | 2.103 | |
| | p | 0.174 | **0.0006** | 0.359 | 0.102 | **0** | 0.0348 | |
| *Zoramia* | z | 2.113 | **5.51** | 2.018 | -0.79 | **-4.373** | **2.976** | 0.984 |
| | p | 0.035 | **0** | 0.041 | 0.251 | **0** | **0.005** | 0.2078 |

**TABLE S2.** Summary of counting parameters used to analyse the topographic density distribution of photoreceptors and ganglion cells (one retina/count).

| **Species** | **SL (mm)** | **PR/GC** | **Grid size (µm)** | **Counting frame (µm)** | **Schaeffer's CE** |
|---|---|---|---|---|---|
| *R. gracilis* | 47.6 | GC | 330x330 | 60x60 | 0.040 |
| *T. fucata* | 60.5 | PR | 610x610 | 100x100 | 0.042 |
| | | GC | 600x600 | 80x80 | 0.037 |
| *O. cyanosoma* | 45 | PR | 550x550 | 80x80 | 0.035 |
| | 45 | GC | 520x520 | 80x80 | 0.045 |
| *O. doederleini* | N/A | PR | 450x450 | 100x100 | 0.036 |
| | 45 | PR | 400x400 | 100x100 | 0.039 |
| *O. notatus* | 50.4 | PR | 550x550 | 80x80 | 0.043 |

PR = Photoreceptors, GC = Ganglion cells, CE = Schaeffer's coefficient of error

**TABLE S4.** Overview of photoreceptor and ganglion cell densities determined in various cardinalfish species. For *T. fucata*, photoreceptor and ganglion cell counts were performed on either eye of the same individual. For *O. cyanosoma*, photoreceptor count and ganglion cell counts are from different individuals.

| Species | Micro-habitat spec | Total number PR | Peak PR cells/mm$^2$ | Total number DC | Peak DC cells/mm$^2$ | Total number SC | Peak SC cells/mm$^2$ | % SC | Total number GC | Peak GC cells/mm$^2$ | Lens ⌀ (mm) | SRP (cpd) |
|---|---|---|---|---|---|---|---|---|---|---|---|---|
| *R. gracilis* | 1 | - | - | - | - | - | - | - | 368,305 | 23,056 | 2.0 | 7.1 |
| *T. fucata* | 3 | 559,756 | 12,300 | 370,176 | 9,500 | 188,781 | 4,700 | 33 | 334,976 | 8,281 | 3.6 | 7.3 |
| *O. cyanosoma* | 3 | 585,384 | 15,468 | 449,262 | 12,188 | 134,219 | 4,063 | 23 | 430,478 | 19,843 | 2.7 | 8.5 |
| *O. doederleini* | 4 | 430,389 | 18,800 | 347,214 | 15,500 | 79,054 | 4,100 | 18 | - | - | - | - |
| | | 414,087 | 18,400 | 327,404 | 14,500 | 79,020 | 4,300 | 19 | - | - | - | - |
| *O. notatus* | - | 616,786 | 16,562 | 449,208 | 12,969 | 167,456 | 4,843 | 27 | - | - | - | - |

DC = double cone, SC = single cones, PR = total photoreceptors, ⌀ = diameter, SRP = spatial resolving power, cpd = cycles per degree

**TABLE S5.** Dataset used for phylogenetic least squares analysis (PGLS). Activity period (N = nocturnal; D = diurnal). Feeding mode (B = benthivore, P+B = benthivore and planktivore, P = planktivore). Microhabitat (specialisation group: 1, 2, 3, 4, 5, 6; based on Figure 2). Proportional opsin gene expression for a given gene relative to total single cone opsin (*SWS2B, SWS2Aα, SWS2Aβ*), total double cone opsin (*RH2B*, *RH2A*, *LWS*), and total opsin (*RH1*) expression (taken from (Luehrmann et al. 2019). N/A = data not available.

| **Species** | | **Predictor variables** | | | **Dependent variables** | | | | | | | |
|---|---|---|---|---|---|---|---|---|---|---|---|---|
| | | | | | | **Proportional opsin gene expression (%)** | | | | | | |
| | | **Activity period** | **Feeding mode** | **Microhabitat partition** | **Relative eye size** | ***SWS2B*** | ***SWS2Aα*** | ***SWS2Aβ*** | ***RH2B*** | ***RH2A*** | ***LWS*** | ***RH1*** |
| *Apogon* | *crassiceps* | N[9,10] | B[3] | N/A | 0.346 | 5.3 | 94.7 | 0 | 0 | 100 | 0 | 98.1 |
| | *melas* | D[12] | N/A | N/A | N/A | 0 | 30 | 70 | 0 | 71.6 | 28.4 | 96.3 |
| *Cheilodipterus* | *artus* | D[12] | P[1] | 3 | 0.492 | 0 | 37.7 | 62.3 | 14.7 | 84.6 | 0.7 | 93.6 |
| | *macrodon* | D[12] | P[1,2] | 4 | 0.491 | 2.3 | 56.1 | 41.7 | 4.4 | 93.4 | 2.2 | 95.1 |
| | *quinquelineatus* | N[10] | B+P[1,2] | 3 | 0.470 | 0.2 | 47.8 | 51.9 | 1.4 | 95.4 | 3.2 | 91.8 |
| *Fibramia* | *thermalis* | D[9] | N/A | 2 | 0.384 | 0.4 | 41.5 | 58.1 | 1.9 | 81.6 | 16.4 | 90.6 |
| *Fowleria* | *variegata* | N[9] | NA | N/A | 0.422 | 0.5 | 35.6 | 64.0 | 0 | 78.2 | 21.8 | 93.1 |
| *Nectamia* | *fusca* | N[3] | P[3] | 6 | 0.527 | 0 | 6.3 | 93.7 | 0.2 | 60.3 | 39.6 | 94.2 |
| | *savayensis* | N[4] | P[3] | 6 | 0.503 | 0 | 8.9 | 91.1 | 0 | 70.6 | 29.4 | 94.0 |
| *Ostorhinchus* | *angustatus* | N[3] | N/A | N/A | N/A | 0 | 23.2 | 76.8 | 0.3 | 99.0 | 0.6 | 91.0 |
| | *compressus* | N[10] | P[5] | 4 | 0.543 | 0 | 38.3 | 61.7 | 7.8 | 92.1 | 0.1 | 90.0 |
| | *cookii* | N[10] | B[6] | 4 | 0.48 | 0.9 | 50.8 | 48.4 | 2.0 | 90.8 | 7.2 | 94.3 |
| | *cyanosoma* | D[12] | B+P[1,2] | 3 | 0.457 | 0 | 24 | 76 | 18 | 81.3 | 0.7 | 92.3 |
| | *doederleini* | N[10] | B+P[1,2] | 4 | 0.469 | 0 | 40.5 | 59.5 | 1.5 | 94.1 | 4.4 | 94.0 |
| | *nigrofasciatus* | N[11] | N/A | 5 | 0.456 | 0 | 65.8 | 34.2 | 3.3 | 96.2 | 0.5 | 91.1 |
| | *notatus* | N[10] | N/A | N/A | 0.479 | 0 | 26.1 | 73.9 | 3.1 | 96.8 | 0 | 92.9 |
| | *novemfasciatus* | N[10] | N/A | N/A | 0.440 | 1.5 | 49.0 | 49.5 | 9.0 | 89.8 | 1.2 | 87.0 |
| *Pristiapogon* | *exostigma* | N[10] | B+P[1,2] | 5 | 0.447 | 0.4 | 99.6 | 0 | 0.1 | 99.9 | 0 | 94.5 |
| | *fraenatus* | N[12] | N/A | N/A | 0.449 | 0 | 100 | 0 | 1.8 | 98.2 | 0 | 92.0 |

| Species | | Predictor variables | | | Dependent variables | | | | | | | |
|---|---|---|---|---|---|---|---|---|---|---|---|---|
| | | | | | | Proportional opsin gene expression (%) | | | | | | |
| | | **Activity period** | **Feeding mode** | **Microhabitat partition** | **Relative eye size** | ***SWS2B*** | ***SWS2Aα*** | ***SWS2Aβ*** | ***RH2B*** | ***RH2A*** | ***LWS*** | ***RH1*** |
| | *cf. mirifica* | N[10] | N/A | N/A | N/A | 0.8 | 22.7 | 76.5 | 0 | 90.1 | 9.9 | 95.9 |
| *Rhabdamia* | *gracilis* | D[4] | P[4] | 1 | 0.389 | 70.1 | 29.9 | 0 | 40.2 | 59.8 | 0 | 64.1 |
| *Sphaeramia* | *nematoptera* | N[7] | P[8] | N/A | 0.494 | 0 | 42.8 | 57.2 | 0.4 | 85.6 | 14.0 | 92.0 |
| *Taeniamia* | *fucata* | N[10] | P[1] | 3 | 0.449 | 0.1 | 17.6 | 82.3 | 1.4 | 96.6 | 2.0 | 91.0 |
| | *zosterophora* | N[10] | P[9] | 3 | 0.470 | 0 | 47.9 | 52.1 | 0.9 | 98.6 | 0.5 | 92.5 |
| *Zoramia* | *leptacantha* | N[10] | P[6] | 3 | 0.469 | 1.8 | 98.2 | 0 | 7.8 | 92.1 | 0.1 | 93.7 |
| | *viridiventer* | N[10] | B+P[1] | 2 | 0.423 | 40.6 | 59.4 | 0 | 13.8 | 86.2 | 0 | 89.8 |

1 - Barnett et al. (2006), 2 - Marnane and Bellwood (2002), 3 - Myers (1999), 4 - Kuiter and Tonozuka (2001), 5 - Job and Shand (2001), 6 - Frédérich et al. (2017), 7 - Lieske et al. (2002), 8 - Nakamura et al. (2003), 9 - Allen et al. (2003), 10 - Paxton et al. (1989), 11 - Randall and Lachner (1986), 12 - Brandl and Bellwood (2014).

**TABLE S3.** PGLS model estimates for correlation of ecological and/or environmental predictor variables and dependent variables. P-values are Bonferroni corrected for repeated hypothesis testing, and statistically significant values are shown in bold. Microhabitat comparison, n = 17 species; Activity period, n = 26 (opsin gene expression), n = 23 (relative eye size); Feeding mode, n = 17; Relative eye size, n = 23.

| Dependent | Predictor | Pagel's λ | $R^2$ | Df | F-Stat | P-value (adj.) |
|---|---|---|---|---|---|---|
| Relative eye size | Microhabitat partition | 0.0 | 0.676 | 5,11 | 7.66 | **0.02*** |
| | Activity period | 1.0 | -0.039 | 1,21 | 0.169 | 0.686 |
| | Feeding mode | 1.0 | 0.278 | 2,14 | 4.078 | 0.322 |
| *SWS2B* | Microhabitat partition | 0.0 | 0.721 | 5,11 | 9.283 | **0.009**** |
| | Activity period | 0.754 | -0.016 | 1,24 | 0.618 | 1 |
| | Feeding mode | 0.896 | -0.128 | 2,14 | 0.093 | 1 |
| | Relative eye size | 0.0 | 0.227 | 1,21 | 7.474 | 0.087 |
| *SWS2Aα* | Microhabitat partition | 1.0 | 0.438 | 5,11 | 3.491 | 0.313 |
| | Activity period | 0.995 | -0.033 | 1,24 | 0.193 | 1 |
| | Feeding mode | 1.0 | -0.044 | 2,14 | 0.664 | 1 |
| | Relative eye size | 1.0 | -0.023 | 1,21 | 0.507 | 1 |
| *SWS2Aβ* | Microhabitat partition | 1.0 | 0.337 | 5,11 | 2.629 | 0.675 |
| | Activity period | 1.0 | -0.041 | 1,24 | 0.011 | 1 |
| | Feeding mode | 1.0 | -0.029 | 2,14 | 0.772 | 1 |
| | Relative eye size | 1.0 | 0.096 | 1,21 | 3.335 | 0.575 |
| *RH2B* | Microhabitat partition | 0.0 | 0.489 | 5,11 | 4.056 | 0.198 |
| | Activity period | 0.776 | 0.133 | 1,24 | 4.842 | 0.264 |
| | Feeding mode | 1.0 | -0.019 | 2,14 | 0.852 | 1 |
| | Relative eye size | 0.816 | -0.043 | 1,21 | 0.100 | 1 |
| *RH2A* | Microhabitat partition | 0.0 | 0.744 | 5,11 | 10.31 | **0.006**** |
| | Activity period | 0.963 | 0.159 | 1,24 | 5.74 | 0.173 |
| | Feeding mode | 1.0 | 0.059 | 2,14 | 1.502 | 1 |
| | Relative eye size | 0.991 | -0.033 | 1,21 | 0.291 | 1 |
| *LWS* | Microhabitat partition | 0.0 | 0.761 | 5,11 | 11.17 | **0.004**** |
| | Activity period | 1.0 | -0.032 | 1,24 | 0.225 | 1 |
| | Feeding mode | 1.0 | -0.133 | 2,14 | 0.061 | 1 |
| | Relative eye size | 1.0 | -0.028 | 1,21 | 0.393 | 1 |
| *RH1* | Microhabitat partition | 0.0 | 0.858 | 5,11 | 20.37 | **<0.001***** |
| | Activity period | 0.941 | -0.042 | 1,24 | 0.000 | 1 |
| | Feeding mode | 1.0 | -0.050 | 2,14 | 0.621 | 1 |
| | Relative eye size | 0.943 | 0.004 | 1,21 | 1.083 | 1 |

Df = degrees of freedom, P-values * < 0.05 , ** < 0.01, *** < 0.001
